# UniGEN-DDI: Computing drug-drug interactions using a unified graph embedding network

**DOI:** 10.1101/2024.09.30.615812

**Authors:** Somnath Mondal, Debarghya Datta, Soumajit Pramanik, Rukmankesh Mehra

**Affiliations:** Department of Chemistry, Indian Institute of Technology Bhilai, Bhilai, Durg-491002, Chhattisgarh, India; Department of Computer Science and Engineering, Indian Institute of Technology Bhilai, Bhilai, Durg- 491002, Chhattisgarh, India; Department of Bioscience and Biomedical Engineering, Indian Institute of Technology Bhilai, Bhilai, Durg-491002, Chhattisgarh, India

**Keywords:** Drug-drug interactions, graph neural network, body part-based split, link prediction, computationally efficient, GraphSAGE, Node2Vec

## Abstract

Finding drug-drug interaction is crucial for patient safety and treatment efficacy. Two drugs may show a synergistic effect but may sometimes cause a severe health issue, including lethality. Wet lab studies are often performed to understand such interactions but are limited by cost and time. However, the biochemical data generated can be explored to compute unknown interactions. Here, we developed a computational model named UniGEN-DDI (<u>Uni</u>fied <u>G</u>raph <u>E</u>mbedding <u>N</u>etwork for <u>D</u>rug-<u>D</u>rug Interaction) for the estimation of interactions between drugs. It is a simple unified network model containing the biochemical information of drug association developed using the compiled data from DrugBank 5.1.0. The feature learning of the drugs was carried out using a combination of GraphSAGE and Node2Vec algorithms, which were found efficient in extracting diverse features. The simple architecture of our model led to a significant reduction in computational time compared to the baselines, while maintaining a high prediction accuracy. The model performed well on the data, which was equally distributed between interacting and non-interacting drugs. As a more challenging evaluation, we performed non-overlapping splitting of the data based on the drug action on different parts of the body, and our model performed well in both interaction estimation and time efficiency. Our model successfully identified the 12 unknown drug interactions in DrugBank 5.1.0, which were updated in DrugBank 6.0.

**Highlights:**

- UniGEN-DDI: A simple unified network model was developed for drug-drug interactions.
- Node embeddings were generated using GraphSAGE and Node2Vec algorithms.
- A significantly higher computational efficiency was achieved compared to baselines.
- Model validated in non-overlapping splitting of data targetting multiple body parts.
- UniGEN-DDI estimated unknown interactions verified in the updated DrugBank 6.0.

**Graphical abstract:** 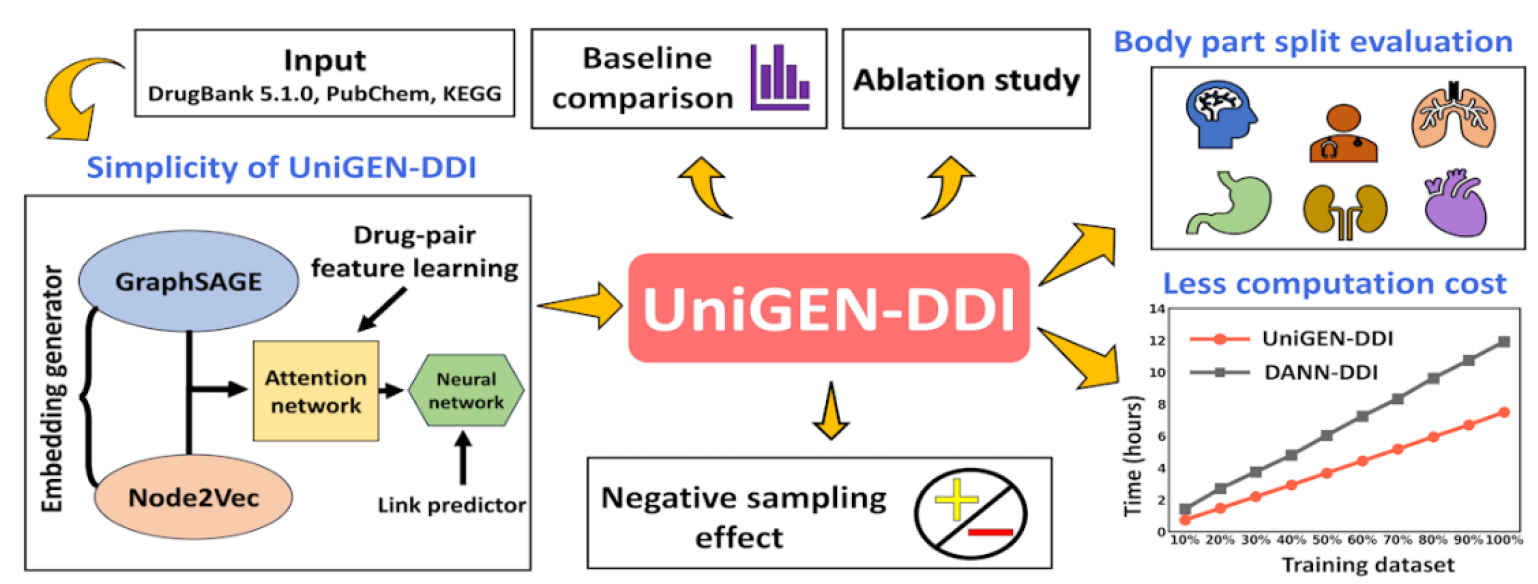

## 1. Introduction

Drug designing is a complex process involving multidisciplinary approaches. Routine drug design does not include the process of evaluating interactions between different drugs. Multiple drugs may be administered to patients to achieve synergistic effects or in case of multimorbidity. Interactions between drugs are commonly studied explicitly after the phase III trials unless a treatment regime involves multiple drugs, which may be tested from the very beginning of preclinical to late clinical trials. Nevertheless, studying the effect of multiple drugs is integral to drug design, but they may only be studied experimentally when two or more drugs are required to be delivered together [1]. Such studies yield essential data on drug interactions commonly available in public repositories such as DrugBank [2] and PubChem [3].

Delivering multiple drugs to patients may cause mild to acute severe effects if administered without the knowledge of possible interactions between them [4,5]. This leads to a crucial question: *whether the drug-drug interactions could be estimated for all the available marketed drugs or the new drugs*. Experimentally, it is a tedious task requiring expensive resources and huge human resources; for example, as of now, there are approximately 1500-2000 marketed drugs, forming nearly about 2 million possible pairs [2]. Experimental evaluation of all possible drug interactions may also not be required unless the need arises, but it is informative to know beforehand in case of urgency. For example, the rapid spread of SARS-CoV-2 infections with a severe mortality rate caused doctors to test several drug combinations within a very short time without much knowledge of the possible interactions between them. Hydroxychloroquine and azithromycin were initially used together in COVID-19 treatment despite concerns about their efficacy and potential cardiac risks [6]. Remdesivir and Dexamethasone later became a common combination for severe cases, as evidence supported their effectiveness [7,8].

Such scenarios revolutionized cheminformatics and structural biology to assist in defining drug combinations. Chemical databases such as DrugBank and PubChem offer insights into medications, while biological databases, including UniProt [9] and PDB [10], provide information on drug-associated proteins. The KEGG [11] database offers knowledge of metabolic pathways. The literature data on the biochemical properties of drugs may be used to develop computational models for estimating new interactions.

The literature studies on drug interactions are discussed in detail in **Section 1** of **Supporting Information**. The DDI prediction techniques can be categorized based on matrix factorization, network, similarity, literature extraction, and deep learning methods [12–16]. In matrix factorization, the DDI matrix can be divided into multiple matrices, which rebuild the interaction matrix to predict DDIs [17–19]. The network-based approaches can directly infer from the network structure or consider the high-order similarity of medications and the transmission of similarity [20–23]. The similarity-based method assumes that medications with similar properties may interact [15,24]. The literature extraction-based method treats the extraction of DDIs as a multi-class classification job, often extracting data from the literature’s instructive phrases before detecting and classifying possible DDIs [25–29]. The deep learning approaches extract high-quality pharmacological characteristics, which have seen extensive use in biology with encouraging outcomes [30–35]. Previously used graph models for predicting DDIs used separate drug-feature networks to build the prediction models, making the prediction task computationally expensive and time-consuming.

Hence, in this work, we aim to propose a DDI prediction framework that not only predicts interactions with high precision but is also computationally efficient. We compiled the data of drug properties from the DrugBank 5.1.0 database and developed a simple unified graph neural network-based model to optimize the time efficacy while maintaining the high estimation ability. This model was named UniGEN-DDI (<u>Uni</u>fied <u>G</u>raph <u>E</u>mbedding <u>N</u>etwork for <u>D</u>rug-<u>D</u>rug Interaction). GraphSAGE [36] and Node2vec [37] algorithms were used in combination with adaptation for link prediction that yielded better performance than using any one of these alone. The negative sampling test containing an equal proportion of interacting (positives) and non-interacting (negatives) samples showed good performance. We adopted a stringent validation approach by non-overlapping splitting of drugs into training and test datasets based on their impacts on various, dividing the dataset based on the drug action on different parts of the body and evaluating testing our model, which performed well.

## 2. Methods

### 2.1 Hypothesis

We aim to build a drug interaction model that could exhibit a high prediction ability with high time efficiency. Computation time could be an important factor when analyzing the interactions aiming at fast predictions as a web tool (especially during critical needs). Since the datasets and models used can have different levels of complexity involved in prediction, building models with low complexity and high accuracy is highly desirable, which could save computational time. To achieve this goal, we built UniGEN-DDI that could compute time-efficient drug-drug interactions with high performance, as further discussed below.

### 2.2 Datasets used

The data from DrugBank released in April 2018 (version 5.1.0), curated by Liu et al. [38], was used. The dataset contains 841 drugs and their known 82,620 pairwise interactions. Important drug features including 619 chemical substructures, 307 metabolic pathways, 214 enzymes, and 1333 targets were considered for expressing the drug’s complete information. The chemical substructures are crucial in determining molecular geometry and physicochemical properties of compounds that impart essential biological functions. Sometimes, a minor change in the chemical structure might transform the drug action [39]. The pathways can provide useful information about the pharmacokinetic actions of the drug [40]. The drugs bind to the target proteins in our body, leading to the remedial effects and forming a crucial biological component of the drug interaction analysis. This provides the mechanistic level information, where a drug may bind to a protein, which may be an enzyme, a receptor, a membrane channel protein or a signaling molecule, etc. [41]. In the DrugBank dataset, the enzyme refers to the protein involved in metabolizing the drug whereas the target is the protein with which the drug binds to show its action. Combining both the chemical and biological characteristics of the drugs may help extract useful biochemical information for building a robust deep-learning drug interaction model.

### 2.3 Graphical representation of drug network

Figure 1. shows the architecture of our model (UniGEN-DDI), containing two modules: (a) drug embedding learning, and (b) drug-drug pairwise feature learning and corresponding interaction estimation. The drug network contained information about drugs, the substructure of drugs, pathways, enzymes, and targets. The concatenation of GraphSAGE and Node2Vec embeddings was used for drug-drug pair feature learning and, finally, generating drug interaction prediction, which are discussed in detail below.

**Figure 1.**
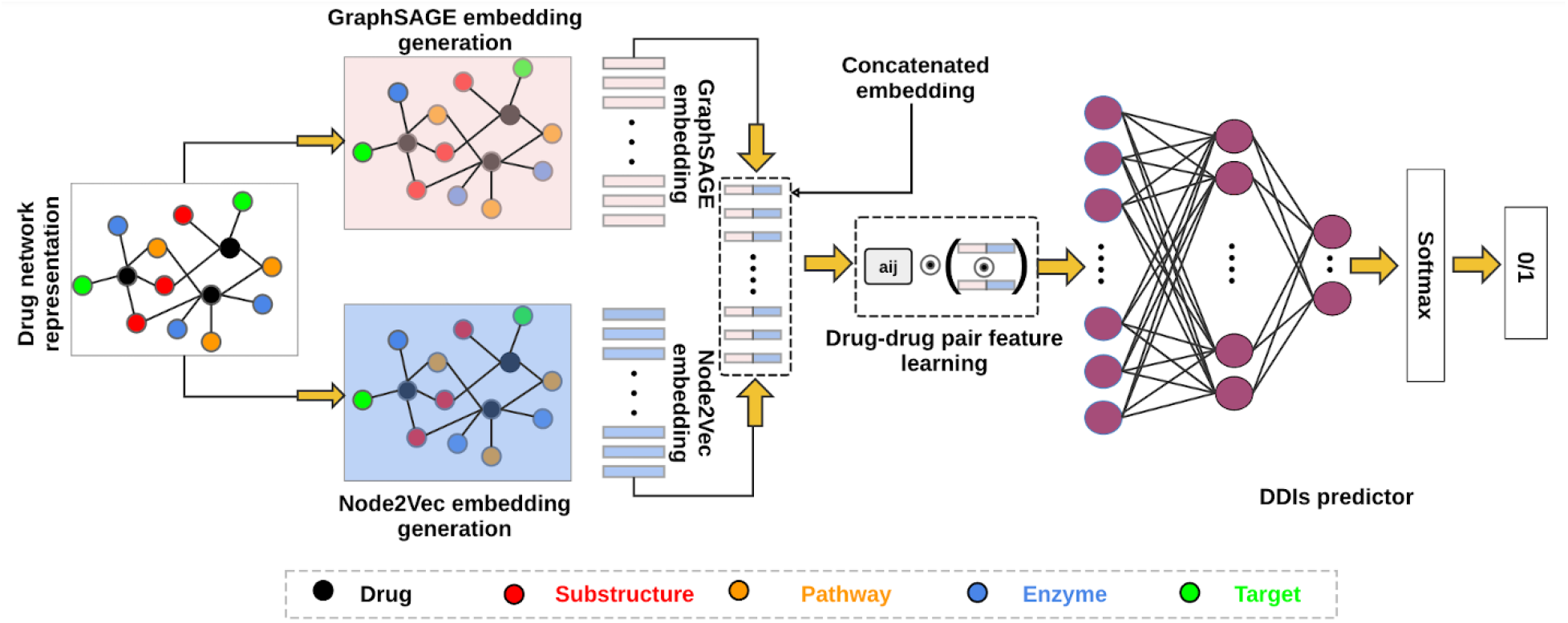
The architecture of our proposed model UniGEN-DDI for drug-to-drug interaction computation.

The drug-drug interaction matrix, *M*, was built, which was a ‘*n x n*’ square matrix, where *n* represents the number of drugs. Each element, *m*_*ij*_, denotes the interaction label between two drugs, *d*_*i*_ and *d*_*j*_. For binary classification, the interaction label is 0 or 1, meaning *m*_*ij*_ *∈* {0,1}, where 0 stands for no interactions, and 1 indicates the existing interactions between two drugs.

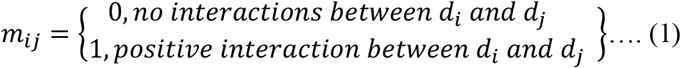

We represented the drug network through graph *G*, where the drugs, pathways (P), enzymes (E), targets (T), and substructures (S), collectively referred to as PETS information, were shown as nodes, and the relations or connections between each node were denoted as edges. The edges were not directional, and no edge weights were assigned. Consequently, the resulting graph network, *G*, encompasses inter-drug relationships and captures the intricate associations between drugs and their pertinent biochemical features. A sample graph is shown in **Figure 2a**.

**Figure 2.**
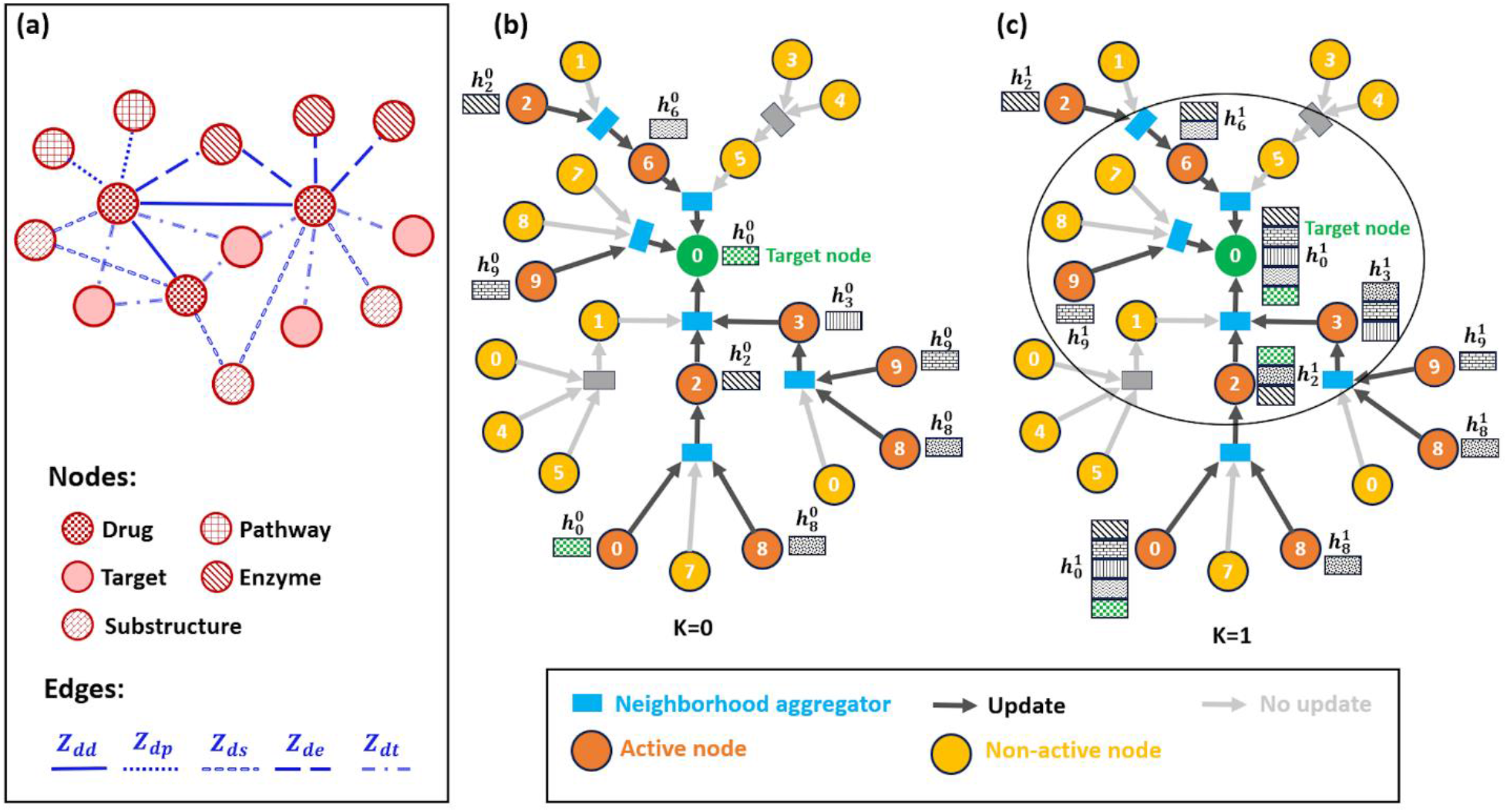
Representation of the graph network and hop aggregation. **(a)** Sample of unified graph network, consisting of drug, pathway, enzyme, target, substructure, and their corresponding association edges (*Z*_*dd*_, *Z*_*de*_, *Z*_*dt*_, *Z*_*dp*_, *Z*_*ds*_). **(b)** Zero hop aggregation representation of GraphSAGE, **(c)** One hop aggregation representation of GraphSAGE.

A graph *G =* (*V, Z*) is defined by a set of vertices *V* and a set of edges *Z*, where *V =* {*V*_*d*_, *V*_*p*_, *V*_*e*_, *V*_*t*_, *V*_*s*_} represents the nodes, and *Z =* {*Z*_*dd*_, *Z*_*de*_, *Z*_*dt*_, *Z*_*dp*_, *Z*_*ds*_} represents the edges. Here, *Z*_*dd*_ *=* {(*v*_*i*_, *v*_*j*_) ∣ *v*_*i*_, *v*_*j*_ *∈ V*_*d*_}, *Z*_*de*_ *=* {(*v*_*i*_, *v*_*j*_) ∣ *v*_*i*_ *∈ V*_*d*_, *v*_*j*_ *∈ V*_*e*_}, *Z*_*dt*_ *=* {(*v*_*i*_, *v*_*j*_) ∣ *v*_*i*_ *∈ V*_*d*_, *v*_*j*_ *∈ V*_*t*_}, *Z*_*dp*_ *=* {(*v*_*i*_, *v*_*j*_) ∣ *v*_*i*_ *∈ V*_*d*_, *v*_*j*_ *∈ V*_*p*_} and *Z*_*ds*_ *=* {(*v*_*i*_, *v*_*j*_) ∣ *v*_*i*_ *∈ V*_*d*_, *v*_*j*_ *∈ V*_*s*_} represent the connections or relationships between different vertices. Each vertex *v*_*i*_ can have attributes denoted by *x*_*vi*_, and each edge (*v*_*i*_, *v*_*j*_) may have an associated weight *w*_*ij*_. The graph’s structure can be characterized using an adjacency matrix *A*, where *A*_*ij*_ = 1 if (*v*_*i*_, *v*_*j*_) in *E* and 0 otherwise. The degree of a vertex *v*_*i*_ is the sum of its adjacent edges, defined as 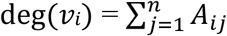.

The graph’s structure can be characterized using an adjacency matrix *A*, where *A*_*ij*_ = 1 if (*v*_*i*_, *v*_*j*_) is in *E* and 0 otherwise. The degree of a vertex *v*_*i*_ is the sum of its adjacent edges, defined as 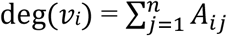.

From the drug-drug interaction matrix *M* and graph network, *G*, our task was to compute whether there were any interactions between *d*_*i*_ and *d*_*j*_, i.e., similar to the link prediction task between two nodes [42]. From equation (1), the value 1 represents the existing interactions between drugs, and 0 indicates no interactions.

### 2.4 Drug feature learning using GraphSAGE

The feature learning of the drugs was performed using GraphSAGE and Node2Vec. GraphSAGE uses a neighborhood aggregation approach for the embedding generation of each node. For the graph *G = (V, E)*, it considers *N(v)* as the neighborhood of node *v*. It initializes the embedding 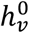 for each node *v* randomly. GraphSAGE aggregates the information from each node at each layer, *l*. If 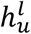 denotes the embedding of neighbor *u* in layer *l*, the aggregated representation 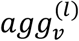 for node *v* is obtained by aggregating the embedding of its neighbors using an aggregation function *AGG*.

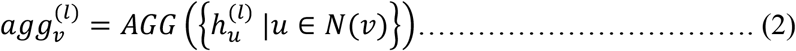

After aggregation, each node’s embedding at layer *l+1* was updated by combining the aggregated representation 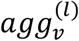 with the current embedding 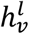 and the node’s own characteristics *x*_*v*_.

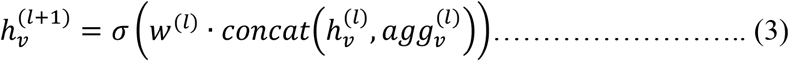

In this equation, *w*^(*l*)^ is the weight matrix of layer *l, σ* is the activation function, and concat denotes concatenation. Thus, aggregation and updating were continued for multiple layers till the convergence or predefined necessary repetitions. After the required number of repetitions, the final embedding *h*_*v*_ for each node *v* was acquired from the GraphSAGE model’s final layer. These embeddings combined both local neighborhood information and the node’s intrinsic properties, and were used for node categorization or connection prediction.

**Figure 2b** and **c**, show zero-hop and one-hop aggregation in GraphSAGE. In zero-hop aggregation, each node’s embedding is generated by directly using its own features without aggregating information from its neighbors. This represents the node’s initial state (can be randomly initialized) or self-information before incorporating any neighborhood information. The one-hop aggregation involves updating the node’s embedding by aggregating information from its one-hop neighbors. The aggregated information from these neighbors is combined with the node’s own embedding to produce an updated representation that reflects both the node’s own features and the influence of its neighbors. Thus, GraphSAGE captures local neighborhood information and (node-specific) features through its aggregation mechanism.

### 2.5 Drug feature learning through Node2Vec

We also used Node2Vec to produce node embeddings. Unlike GraphSAGE, which aggregates information from node neighborhoods, Node2Vec learns embeddings by performing random walks on the graph and training a Skip-gram model to predict surrounding nodes based on a target node [43]. The learned embeddings capture the structural features of the graph’s nodes.

For each node *v* in the graph *G =* (*V, E*), it performs *r* random walks of length *l*, resulting in sequences *S*_*v*_ *=* {*v, u*_1_, *u*_2_, …, *u*_*l*_}, where *u*_*i*_ are the nodes visited during the walk. For a given target node *v* in a sequence *S*_*v*_, it maximizes the probability of observing its neighboring nodes *u*_*i*_ in the context window. The likelihood of observing a neighboring node *u*_*i*_ for given the target node *v* is given by the equation:

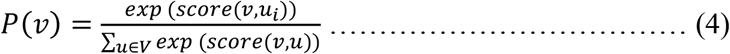

The scoring function *score*(*v, u*) measures the similarity between the embeddings of nodes *v* and *u*. The dot product or cosine similarity is commonly used as a scoring function. We gave a random initialization to the node embeddings.

The objective function of the Skip-gram model is to maximize the log-likelihood: *max ∑*_*v∈V*_ *∑*_*ui∈Sv*_ *P*(*u*_*i*_|*v*). Every node *v* is connected to a low-dimensional embedding vector *z*_*v*_ that is generated during the training phase of the Skip-gram model. These embeddings encode details about the node’s local and global connectivity patterns in the graph, capturing their structural characteristics.

### 2.6 Loss function

For both GraphSAGE and Node2Vec, we employed the binary cross-entropy loss function with logits, defined as:

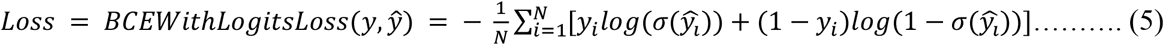

Here *y* is the ground truth label, ŷ the predicted score, and *σ* the sigmoid function. BCEWithLogitsLoss is used to backpropagate and train the networks by calculating the loss between the predicted output and the ground truth labels, which is then used to update the network’s weights during training through backpropagation.

### 2.7 Drug-pair feature learning and Drug-drug interaction estimation

Prior research on attention networks has shown promise in node representation learning scenarios [44–48]. To learn representations of drug pairs, we fused learned drug representations from drug feature networks to create the drug pair feature learning component, which is an attention-based neural network.

We concatenated GraphSAGE and Node2Vec embeddings and get a combined representative embedding 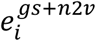 for drug *d*_*i*_, *i =* {1,2, …, *n*}. Then, we introduced the K-dimensional attention vector (*a*_*i,j*_) in such a way that the drug-drug representation of *d*_*i*_ and *d*_*j*_ was the element-wise product among 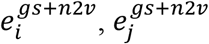, and *a*_*i,j*_.

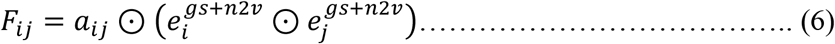

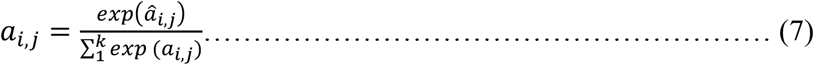

Here, 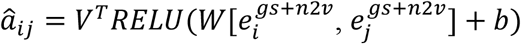, where *RELU* is an activation function. *W* is the weight matrix, *b* is the bias vector, *V*^*T*^is the weight vector, and 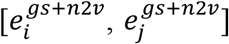 is the concatenated embeddings of *d*_*i*_ *and d*_*j*_. Attention mechanisms are employed to dynamically weight the contributions of different features in the combined embeddings. It allows the model to focus on the most relevant parts of the embeddings for predicting the DDIs, ensuring that higher weights are given to the important features. Thus, by emphasizing the relevant features and de-emphasizing irrelevant ones, the attention mechanism leads the model to better performance in DDIs prediction task.

The main components of our drug-drug interaction estimation comprised the input layer, the fully connected hidden layers, and the output layer. Two neurons in the output layer may indicate interaction or non-interaction, whereas the input layer has the same dimension as the drug-drug pair representation. We employed *RELU* [49] as the activation function for all hidden layers and used the *SoftMax* [50] function in the output layer to generate probabilities. We used binary cross-entropy as a loss function and utilized the Adam optimizer with its default parameters to optimize drug-drug interaction prediction.

### 2.8 Parameters optimization

We performed model optimization by varying various parameters. Our proposed model has two drug-feature learning components: (a) For GraphSAGE, we conducted a grid search for all sets of parameters and found the best performance using two layers, batch size of 256, dropout of 0.5 in each hidden layer, a learning rate of 0.001 and 150 epochs for the best performance. (b) For the Node2Vec embedding generator, we used a walk length of 40, 10 walks per node, a context size of 20, a 256-dimensional batch size, a learning rate of 0.01, and 150 epochs. These are discussed in the Results section.

For the drug-drug interaction component, we used seven fully connected hidden layers with a dropout rate of 0.4 for each, 150 epochs, and an early stop to avoid overfitting the model.

### 2.9 Comparisons to baselines

To evaluate the performance, we compared our model with four other DDI prediction methods, including DANN-DDI [38], DDI-MDAE [30], DPDDI [14], and RANEDDI [51]. The DANN-DDI method used a graph-based learning approach to obtain drug embeddings from four different drug feature networks. The learned drug embeddings were concatenated, and an attention neural network was proposed to learn representations of drug pairs. A deep neural network was employed to predict drug-drug interactions and their associated events.

The DDI-MDAE method is a multi-modal deep auto-encoder-based drug representation learning method, which incorporates the information about interactions, substructures, targets, enzymes, and transporters of the drugs. DPDDI comprised a graph convolutional network-based feature extractor for drug graph embeddings and a deep neural network-based predictor concatenating drug latent feature vectors to predict potential interactions. The RANEDDI method combined relation-aware network structure information into the topology of a multi-relational DDI network to embed medicines. This model used relationship-aware network embedding to predict drug interactions.

### 2.10 Primary evaluation metrics

The performance of the model was primarily evaluated using the standard metrics for binary classification prediction, including recall, precision, F1 score, and accuracy (ACC) as shown below in **equations 8-11**:

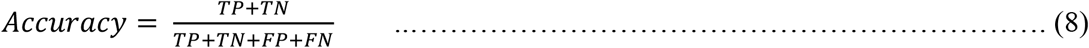

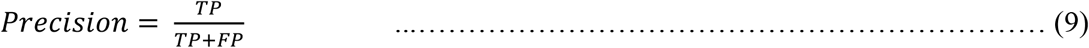

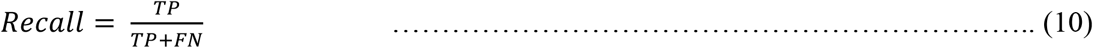

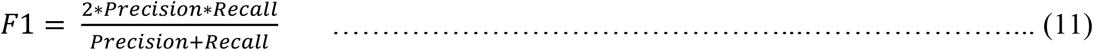

TP represents correctly identified drug-drug interactions. TN indicates accurately identified absences of drug-drug interactions. FP are false positives, which falsely identify the presence of interactions between drugs that are actually not present. FN refers to falsely identified absences of a drug-drug interaction that is actually present. Accuracy is the percentage of accurately detected interactions (including true positives and true negatives) among all drug pairs. Precision is the proportion of true positive identifications to total positive identifications. Recall is the proportion of true positive identifications among all actual positives. F1 score is the harmonic mean of precision and recall, which balances both measurements.

### 2.11 Secondary evaluations

After building and optimizing the model using parameter variation and primary analysis, we performed the validation of our model using multiple stringent criteria. First, we randomly assembled negative samples from the complement set of positive samples, ensuring an equal balance of positive and negative samples throughout all process phases. We employed a random division strategy to allocate approved DDIs into training, validation, and testing sets, maintaining an 8:1:1 ratio across all sets. We used binary cross-entropy as the loss function and the Adam optimizer for optimization [52,53]. Additionally, we inserted batch normalization layers between hidden layers to speed up convergence and include dropout layers to mitigate overfitting and enhance generalization.

Second, we introduced splitting the data based on the drug’s action on different body parts and testing our model’s performance, evaluating tough and more generalized limits [54]. The data were divided into twelve groups of the drugs acting on: (1) circulatory system, (2) digestive system, (3) endocrine system, (4) immune system, (5) integumentary system, (6) lymphatic system, (7) musculoskeletal system, (8) nervous system, (9) reproductive system, (10) respiratory system, (11) urinary system, and (12) the drugs that were associated with more than one type of body part were categorized as whole body part. The details of the dataset division are shown in **Tables S1** and **S2**. In this case, we divided the data into the training and test sets in such a way that the test set contains one of the splits, and the training set includes the rest of them. This ensures that during training, we have no information about the drugs we would find in the test set, making it a more challenging problem. We maintained the same number of training edges (by sampling) to fairly evaluate each body split.

## 3. Results and discussion

### 3.1 Link prediction and baseline comparison

In UniGEN-DDI, the drug information was represented in a single graph network, where a complete drug association network, including pathways, enzymes, targets, and substructures, was presented. We used GraphSAGE and Node2Vec for drug feature learning tasks. GraphSAGE promptly incorporates graph structure, captures local topology through aggregation, scales to huge graphs, and offers configurable embeddings that may be used for various graph types. It permits node information to be included, improving the model’s prediction performance. On the other hand, Node2Vec embeddings perform well in link prediction by rapidly capturing local and global structural information in networks using the node2vec technique, increasing predictive accuracy. We mainly concatenated the GraphSAGE and Node2Vec embeddings for the drug feature learning task. Combining GraphSAGE’s localized neighborhood-aware embeddings with Node2Vec’s globally informed embeddings resulted in a nuanced blend representation of local and global graph structure. This complete embedding improved comprehension of node interactions and led to a more accurate predictive model. The combination took advantage of the complementary nature of the two approaches, with GraphSAGE focusing on capturing local topology and node2vec focusing on balancing local and global exploration.

**Table 1** shows the comparison of our model with the earlier proposed baselines. Interestingly, UniGEN-DDI showed better prediction than DDI-MDAE, DPDDI, and RANEDDI and nearly similar results to DANN-DDI (**Figure 3a**). DDI-MDAE seems to show the lowest values, whereas UniGEN-DDI and DANN-DDI showed comparable results with the highest values of 0.99 for the parameters shown in **Table 1**. It is imperative to note that the highest possible value for these parameters representing performance is 1.00 and it is highly unlikely to achieve values >0.99, unless for a very typical case. So, 0.99 could be considered the best possible score for these parameters if the models perform well. As mentioned, in our case, both UniGEN-DDI and DANN-DDI are high-performing models. In UniGEN-DDI, we captured the complete information of drug-drug and other drug association interactions using a unified drug network and generating embedding with the help of GraphSAGE and Node2Vec model. In contrast, DANN-DDI used five different graphs through a structural deep network embedding (SDNE) and an attention-based neural network.

**Table 1.** Comparison of our model with the baseline models. The values in bold are the best reported and the underlined values are the second best.

| **Model** | **Precision** | **Recall** | **F1** | **Accuracy** |
| --- | --- | --- | --- | --- |
| UniGEN-DDI | **0.99** | **0.99** | **0.99** | **0.99** |
| DANN-DDI | **0.99** | **0.99** | **0.99** | **0.99** |
| DDI-MDAE | **0.99** | 0.68 | 0.80 | 0.97 |
| DPDDI | 0.96 | <u>0.98</u> | <u>0.97</u> | 0.97 |
| RANEDDI | **0.99** | 0.96 | <u>0.97</u> | <u>0.98</u> |

**Figure 3.**
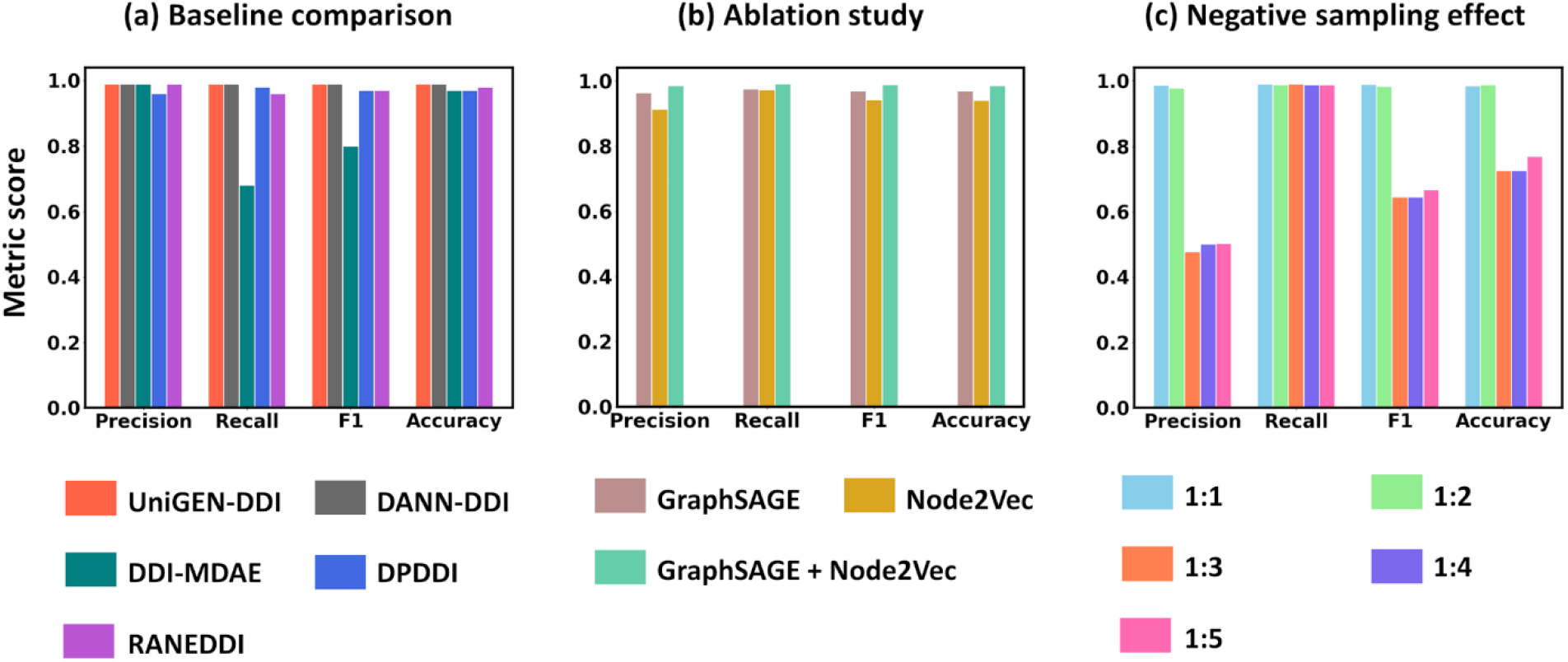
Statistical analysis of our UniGEN-DDI model. **(a)** Baseline comparison with other models. **(b)** Ablation test. **(c)** Negative sampling test using positive:negative samples of 1:1 (sr1 dataset), 1:2 (sr2), 1:3 (sr3), 1:4 (sr4), and 1:5 (sr5).

Now, the next possible improvement could be to evaluate the complexity and therefore, the models’ computational cost, where UniGEN-DDI clearly outperformed DANN-DDI, as discussed later in this work.

### 3.2 Ablation study to understand the contribution of embeddings

To understand the contribution of each component to the overall performance of UniGEN-DDI, we performed the ablation study on the dataset containing the positive to negative sample ratio of 1:1 (named sr1) by random selection. The unlabeled drug combinations were assigned negative samples. For this, three different experimental setups were created using the following embeddings: (a) only GraphSAGE, (b) only Node2Vec, and (c) the GraphSAGE and Node2Vec together. All three setups showed a decent performance (**Table 2** and **Figure 3b**); however, the best was achieved when GraphSAGE and Node2Vec embeddings were combined, indicating that both were essential.

**Table 2.** Ablation study using three different embedding sets. The values in bold are the best reported and the underlined values are the second best.

| Metrics | Precision | Recall | F1 | Accuracy |
| --- | --- | --- | --- | --- |
| GraphSAGE | <u>0.96</u> | <u>0.98</u> | <u>0.97</u> | <u>0.97</u> |
| Node2Vec | 0.91 | 0.97 | 0.94 | 0.94 |
| GraphSAGE + Node2Vec | **0.99** | **0.99** | **0.99** | **0.99** |

### 3.3 Effect of negative sample size on our model

Balancing a dataset ensures that the model does not become biased towards the majority class, thus improving its ability to recognize and correctly classify the minority class. This leads to more accurate and meaningful performance metrics [55]. To address the imbalance in our dataset, which contains more negative interactions than positive ones, we created a negative sample set by sampling various unlabeled drug combinations. Combining these negative samples with known drug-drug interactions (DDIs) pairs generated training, validation, and test datasets with different positive-to-negative sample ratios: 1:1, 1:2, 1:3, 1:4, and 1:5. These ratios, denoted as sr1, sr2, sr3, sr4, and sr5 respectively, were used to assess how varying the proportion of negative samples influences the model’s performance.

**Figure 3c** and **Table S3** depict the findings of our model on these five datasets in the five cross-validation tests. Clearly, the sr1 and sr2 datasets with 1:1 and 1:2 of positive: negative sampling showed high performances in all aspects. However, the overall trend of the metrics indicates that sr1 performed slightly better than sr2. These observations suggest that our model was able to consider the impact of negative data sampling when either an equal number or twice the negative number of samples were considered in the datasets. This makes absolute sense because the dataset having equal representation of the positive and negative sets using random sampling provides the best possible measure of the model performance. An increase in the number of negative sample sets reduced the performance, as obvious, because then the dataset was biased towards the large distribution of the negative set, which was undesirable.

### 3.4 Effect of GraphSAGE layers, batch size, and learning rate

**Figure 4** shows the effect of the number of GraphSAGE layers, batch size, and learning rate on our model’s precision, accuracy, F1, and recall. The GraphSAGE was studied by varying the number of layers using a set of {1,2,3,4,5} layers, keeping all the other parameters constant. Interestingly, our model UniGEN-DDI performed well with two GraphSAGE layers (**Table S4**). This indicates that a single layer in GraphSAGE may not capture enough complexity or depth in the data, resulting in underfitting, and it may lack the capacity to learn the intricate patterns present in the data, resulting in lower performance than two layers, which seems to provide a better representation of the underlying relationships. Adding a layer reduced the score that may be due to the unnecessarily increased complexity or overfitting of the data.

The analysis of the batch size dimensions of {16, 32, 64, 128, 256, 512} in GraphSAGE showed an optimal result at 256-dimensional size (**Figure 4b** and **Table S5**). The learning rate indicates the rate or the speed at which the model learns. Its lower value shows a higher learning rate. In each hidden layer of GraphSAGE, our model outperformed other known methods at the optimal learning rate of 0.001 (**Figure 4c** and **Table S6**), which is well within the acceptable range and therefore, indicates the higher speed and lower computation time at which the model learns.

**Figure 4.**
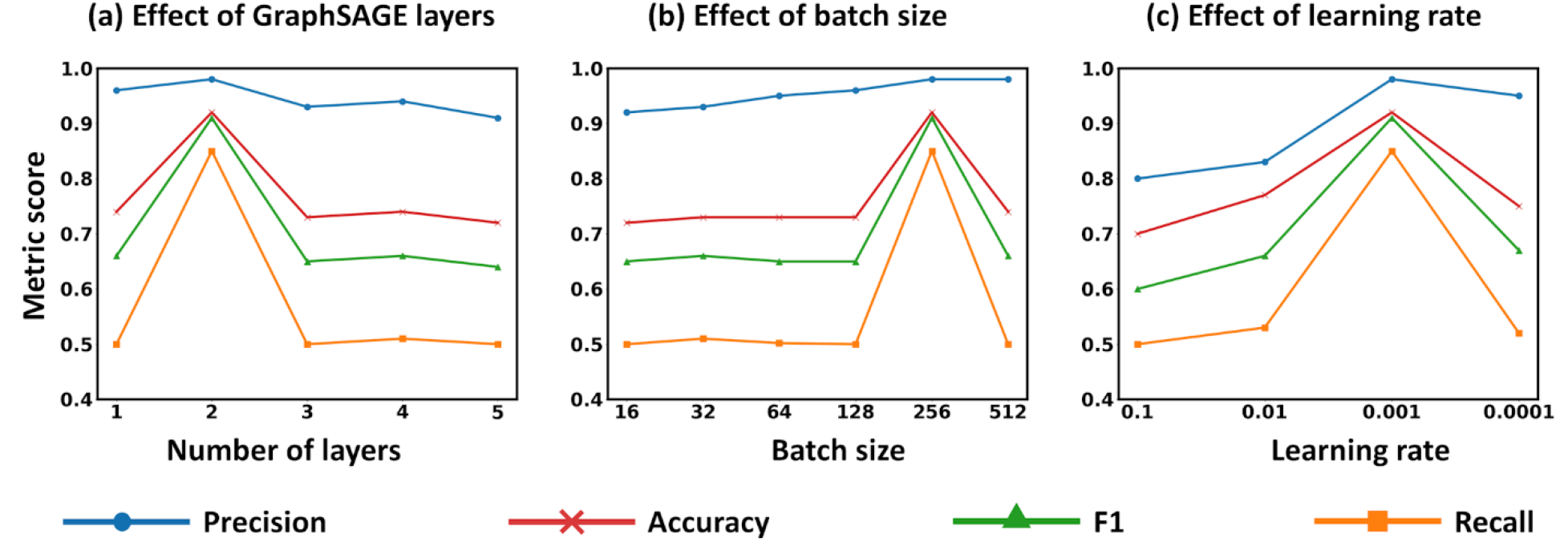
Effect of varying parameters in GraphSAGE on our UniGEN-DDI model. **(a)** Varying GraphSAGE layers using a set of {1,2,3,4,5} layers. **(b)** Performance at different batch sizes of {16, 32, 64, 128, 256, 512} dimensions. **(c)** Understanding the optimal learning rate for our model’s best performance using grid search of learning rate from the set {0.1, 0.01, 0.001, 0.001}.

### 3.5 Higher computational efficiency of our model owing to its simple architecture

As discussed above, both UniGEN-DDI and DANN-DDI showed a similar evaluation of the metric parameters (**Figure 3a** and **Table 1**). However, UniGEN-DDI performed significantly better than DANN-DDI in terms of time efficiency. To analyze this, we computed the training time for both these models using the best-performing dataset sr1 (Figure 3c and Table S3), which contained a 1:1 positive to negative sample ratio.

This analysis began with computing the training time while keeping 10% of the total edges randomly to train the model and then gradually increasing the number of edges by 10% randomly till 100% of the edges were included in the training set. **Figure 5** and **Table S7** compare the training time of the two models. Clearly, our model UniGEN-DDI performed better than DANN-DDI using all training sets, requiring comparatively less training time. Importantly, as we increased the number of edges in training, the differences in time became more dominant.

**Figure 5.**
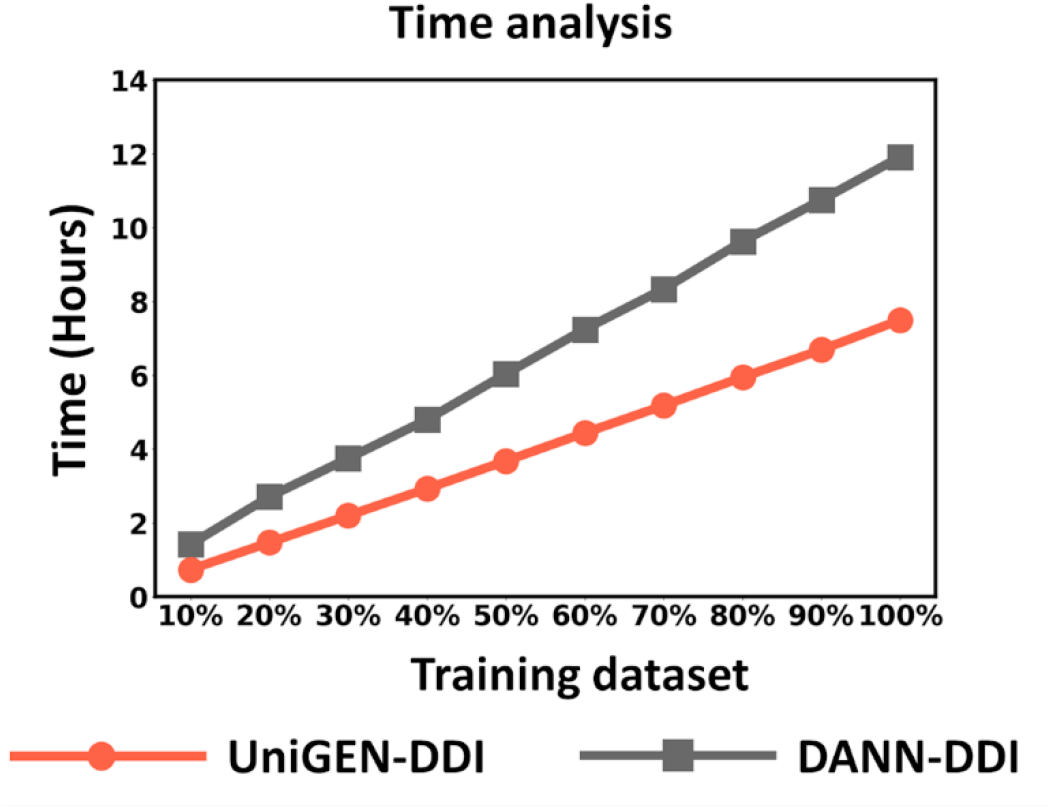
Comparison of the training time between UniGEN-DDI and DANN-DDI. The training dataset is shown in the X-axis starting from 10% of the edges with an increment of 10% up to 100%. The Y-axis represents the computing time on the sr1 dataset.

We further compared the model complexity of DANN-DDI with our model in terms of architecture, which is related to computational efficiency. DANN-DDI was trained using five different graphs for the drug-feature learning that employed the SDNE algorithm [56]. This appears to be a complicated model that, though exhibiting good performance, required high computational power. On the other hand, our model UniGEN-DDI, which exhibited a good performance at the level of DANN-DDI, was based on a simple architecture using GraphSAGE and Node2Vec and generated drug embedding using a single graph network. The simplicity of our approach made our model capable of reducing the computational cost as DANN-DDI needs to retrain all five networks in case new nodes/interactions are added, whereas we need to train only one.

### 3.6 Performance of our model in different body parts

As discussed above, the simple architecture of our model UniGEN-DDI, containing a single graph network, performed well in metric evaluation, baseline comparisons, negative sampling, and most importantly, in computing time. We wanted to check the rigid limits of our model and therefore, adapted a strategy to split the dataset into twelve groups (as described in Methods) based on the effect on various body parts and test the prediction ability of our model. This led to a hard and robust prediction task for the model, also because one of the twelve groups was kept in the test and the ten in the training set. One group containing the lowest number of data points was excluded from the analysis.

Notably, our model showed better performance than baselines DDI-MDAE, DPDDI, and RANEDDI, and was comparable to DANN-DDI (**Figure 6**), which was similar to above baseline comparisons. Here again, our model required less time for computation than DANN-DDI, which is a very crucial factor when dealing with a large dataset, putting our model a step ahead. The detailed results are shown in **Figure S1** and **Table S8**, suggesting that UniGEN-DDI generally outperformed the baseline methods in terms of both metric score and computational time. This showed the robustness of our model, which can be generalized to perform non-trivial estimates without relying solely on the previously known interactions for training.

**Figure 6.**
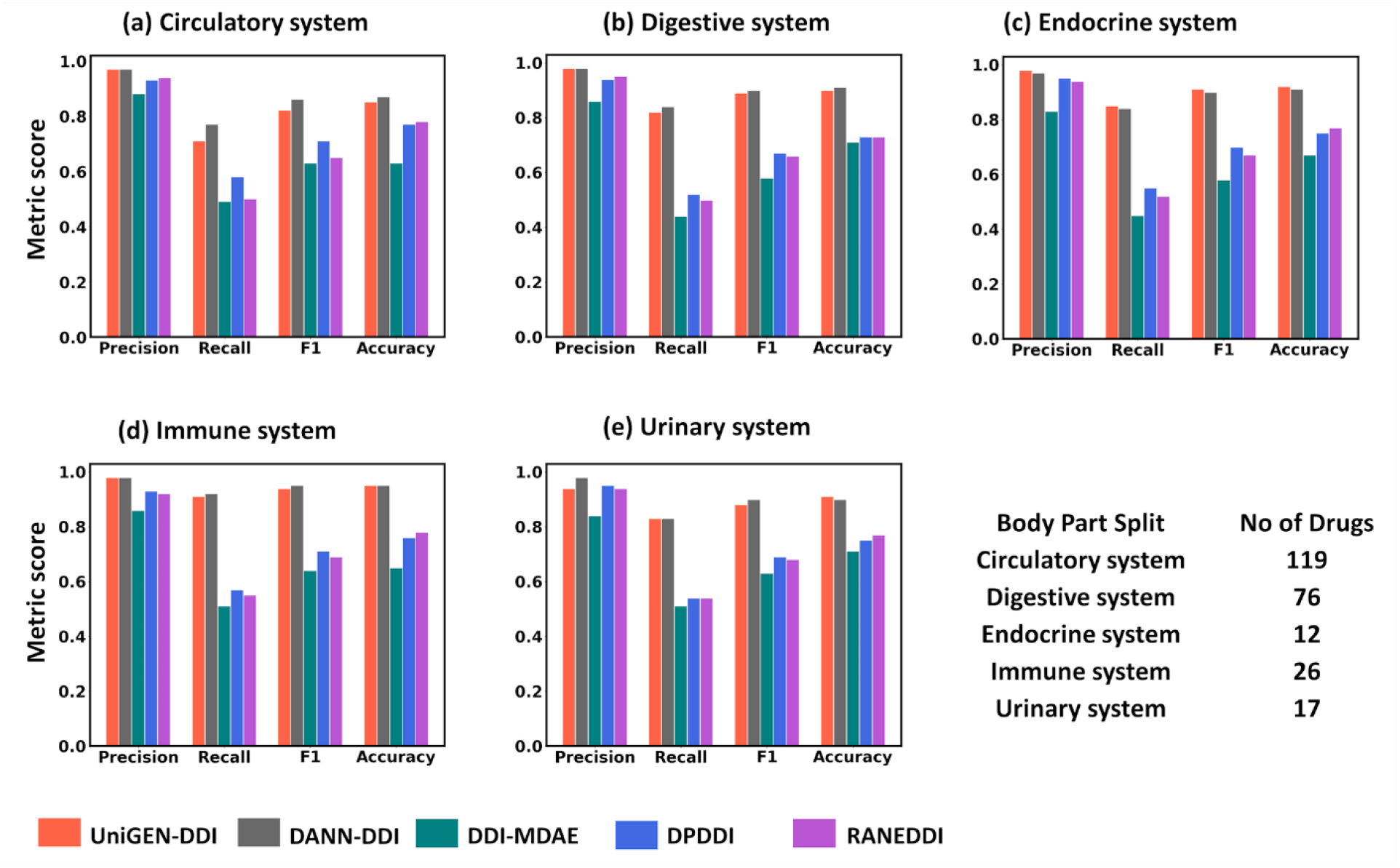
Comparison of UniGEN-DDI with baseline models in different body part splits. **(a)** Circulatory system. **(b)** Digestive system. **(c)** Endocrine system. **(d)** Immune system. **(e)** Urinary system.

### 3.7 Verification of the estimated drug interactions

UniGEN-DDI predicted drug-drug interactions based on DrugBank (version 5.1.0) dataset. However, by the time of the current model completion, DrugBank (version 6.0) was released. Now, it was essential to see whether the interactions predicted by UniGEN-DDI appeared in the updated version of DrugBank or not. This would help define the robustness of the model.

In this regard, we analyzed drug-drug interactions estimated by our model for the presence in Drug 6.0, as shown in **Table 3**. Our model effectively predicted twelve recent case studies sourced from various references in DrugBank 6.0 [57].

**Table 3.** Case studies based on DrugBank 6.0.

| **Drug 1** | **DrugBank Id** | **Drug 2** | **DrugBank Id** | **Evidence from DrugBank 6.0** |
| --- | --- | --- | --- | --- |
| Alfuzosin | DB00346 | Thiothixene | DB01623 | Increases the risk or severity of QTc prolongation. |
| Alfentanil | DB00802 | Bumetanide | DB00887 | Likely to decrease the therapeutic efficacy of bumetanide. |
| Ambrisentan | DB06403 | Celiprolol | DB04846 | Celiprolol has the potential to enhance the hypotensive effects of ambrisentan. |
| Bisoprolol | DB00612 | Lidocaine | DB00281 | May increase the serum concentration of lidocaine. |
| Cinchocaine | DB00527 | Doxepin | DB01142 | Potentially increases the risk or severity of CNS depression. |
| Maraviroc | DB04835 | Ivacaftor | DB08820 | Possibility to increase the serum concentration of maraviroc. |
| Lumacaftor | DB09280 | Treprostinil | DB00374 | Potentially decrease the metabolism of treprostinil. |
| Pindolol | DB00960 | Vortioxetine | DB09068 | Likely increase the risk or severity of CNS depression. |
| Hydromorphone | DB00327 | Imidafenacin | DB09262 | Increases the risk of adverse effects, e.g. urinary retention, paralytic ileus, etc. |
| Vemurafenib | DB08881 | Dexamethasone | DB01234 | Possibility to increase the metabolism of vemurafenib. |
| Lubiprostone | DB01046 | Umeclidinium | DB09076 | Lubiprostone can potentially reduce the excretion rate of vilanterol, leading to a possible increase in serum levels. |
| Albendazole | DB00518 | Bosentan | DB00559 | Potentially increase the metabolism of albendazole. |

As further updates would be available in the DrugBank data, with wet lab experimental studies confirming the drug interactions, the model could be made more robust. In the context of drug-drug interactions, a score of 1 denotes a confirmed relationship between two medications. In contrast, a score of 0 signifies an uncertain association, which doesn’t necessarily imply no interaction. These instances labeled as 0 could potentially be reclassified as 1 in the future if confirmed in *in vitro* or *in vivo* experiments, indicating a verified relationship. The purpose of including these uncertain drug-drug connections in this section is to highlight the efficacy of the proposed framework in our experiments.

## 4. Conclusions

Computing drug-to-drug interaction is an essential task highlighting the potential of multiple drugs to be delivered together in case of acute illness. Predicting beforehand the possibility of ill effects caused by interactions may help in a better treatment regime. It seems to be a difficult job since it highly depends on the type and extent of data availability. Data scientists are working to find these interactions, but the limitation is the non-availability of sizable, refined datasets and the dataset distributions of the positive and negative datasets. We mostly have the data that says that the drugs are interacting, but we hardly find confirmed data that suggest the two drugs do not have interactions. This might be because wet lab experiments are often performed to understand the effect of two or more drugs required in multimorbidity conditions, but not for all possible drug combinations available in the market. This means that all potential drug interactions are not known. Here, computational studies play a vital role in extracting patterns from the available data and computing unknown drug interactions.

Therefore, in the present study, we compiled the dataset of drug interactions from DrugBank (5.1.0) and built a simple single network model using GraphSAGE and Node2Vec for drug feature learning. A complete drug association network containing information on pathways, enzymes, targets, and substructures was presented. Notably, our model outperformed the commonly known baselines and showed comparable results to the DANN-DDI model. The ablation study showed the necessity for both GraphSAGE and Node2Vec embeddings for complete information on drug networks. Our model performed well for the data containing an equal proportion of positive and negative samples.

Computing time for the training set could be an essential feature for any prediction tool as many a time it has to deal with large datasets. Our model UniGEN-DDI clearly performed all the baseline significantly, and the differences in time became prevalent with increasing training data. This is an important outcome which makes any model viable. The time efficiency could be related to the simple architecture of our model, which used a single unified graph network to present drug associations, whereas DANN-DDI considered five different networks to present each drug-pathway, drug-enzyme, drug-target, drug-substructure, and drug-drug associations, which makes it computationally expensive.

To check the hard limits of our model, we performed the non-overlapping splitting of the data based on the drugs acting on different body parts. We introduced this new strategy to evaluate the model’s performance, which consistently showed that UniGEN-DDI performed better than the baselines when considering metric and time performances.

Toward the end of this study, we found that the DrugBank 6.0 version was released. So, it became imperative to test the predictions of our model for the occurrence of any newly added interactions absent in the earlier versions. Notably, twelve drug interactions estimated by our model were defined in the updated DrugBank version. All these observations suggested our model’s robustness and time efficiency, which would be helpful in future interaction predictions, and the model can be further improved with more data availability.

## Supporting Information

Details about the work are provided in the supporting file “supplementary.pdf.”

## CRediT authorship contribution statement

### Rukmankesh Mehra

Conceptualization, Data curation, Formal analysis, Funding acquisition, Investigation, Methodology, Project administration, Resources, Software, Supervision, Validation, Visualization, Writing – original draft, Writing – review & editing. **Somnath Mondal:** Data curation, Formal analysis, Investigation, Software, Validation, Visualization, Writing – original draft, Writing – review & editing. **Debarghya Datta:** Data curation, Software, Validation.

### Soumajit Pramanik

Methodology, Software, Supervision, Validation, Writing – review & editing.

## Acknowledgments

RM and SM acknowledge SERB-SRG, Govt of India, for funding via grant SRG/2022/000304. RM acknowledges IIT Bhilai for supporting this work via the Research Initiation Grant (RIG), number 2005900. SM acknowledges the Department of Science and Technology, Govt of India, for providing the funds to carry out the research under file no. DST/INSPIRE Fellowship/2023/IF230110.

