## Supplementary Information for "UniGEN-DDI: Computing drug-drug interactions using a unified graph embedding network"

### Section 1. Literature review

Drug-drug interaction (DDI) occurs when two or more drugs interact, affecting their competence, potency, or tolerability and potentially amplifying or diminishing their intended effects, leading to unexpected consequences and significant harm to the user [1,2]. DDIs arise from various factors, including pharmacokinetics and pharmacodynamics, patient characteristics, dosage, frequency of administration, and unique mechanisms of action of the involved pharmaceuticals. Pharmacokinetic interactions, for instance, involve changes in medication absorption, distribution, metabolism, or excretion. This can alter the presence of medicine in the blood levels, potentially leading to toxicity or reduced efficacy [3]. On the other hand, pharmacodynamic interactions entail a drug's direct impact on another drug's intended therapeutic or harmful effects, often via shared or opposing pathways in the body [1].

Healthcare professionals play a crucial role in recognizing and managing DDIs by carefully assessing a patient's medical history and current prescription regimen. Understanding these interactions is paramount for optimizing treatment regimens and ensuring patient safety [4,5]. To comprehend DDIs, one must grasp the fundamentals of medications, including their characteristics, structure, metabolism, intended targets, and associated pathways. Information from both cheminformatics and bioinformatics is essential in this regard [6,7].

Computational models are employed to examine the chemical characteristics of medications and their potential interactions with biological entities, and by foreseeing and understanding these interactions, researchers can anticipate potential adverse effects when multiple medications are taken concurrently [7]. Clinical trials are essential for evaluating drug efficacy and safety, but they have limitations such as time constraints, cost, and practical constraints [8]. To address these challenges, researchers have turned to computational approaches for predicting DDIs. Current computational DDI prediction techniques can be categorized into matrix factorization-based, network-based, similarity-based, literature-extraction-based, and deep learning-based methods [9–14].

The drug-drug interaction matrix is divided into various matrices using matrix factorization-based algorithms, which rebuild the interaction matrix to forecast DDIs. Shi et al. used the triple matrix factorization model to predict comprehensive DDIs and binary DDIs [15]. Zhang et al. established a novel matrix factorization approach called MRMF and introduced manifold regularization based on drug characteristics [16]. For enhancive and degressive DDI prediction, Yu et al. developed DDINMF based on semi-nonnegative matrix factorization, which decomposes the DDI adjacent matrix by the regular nonnegative matrix factorization (NMF) [17].

Network-based approaches can directly infer from the network structure or consider the high-order similarity of medications and the transmission of similarity. A label propagation approach was presented by Zhang et al. that focused on high-order similarity and calculated the similarities of side effects, off-label side effects, and chemical structure [18]. To mimic signaling transmission on the protein-protein interaction (PPI) network, Park et al. employed a random walk technique.[19] To uncover the routes of drug-drug couples, Lee et al. built a heterogeneous bioinformatics network and used graph traversal methods [20]. Huang et al.

established the S-score metric to assess the strength of network connections and used the Bayesian probability model to provide precise prediction outcomes [21].

Methods based on similarity assume that medications with similar properties may interact. Gottlieb et al. created the INDI model to predict drug-drug interactions, which computed seven categories of similarity and utilized a weighted logistic regression classifier [22]. To define drug-drug pairings, Cheng et al. included a range of drug-drug commonalities and built prediction models using five classifiers [23].

Literature extraction-based approaches treat the extraction of DDIs as a multi-class classification job, often extracting data from literature's instructive phrases before detecting and classifying possible DDIs. Sun et al. built a convolutional neural network-based deep architecture with several layers of tiny convolutions to automatically extract information [24]. Lexical and syntactic characteristics were incorporated into a linear kernel-based classifier by Kim et al. to increase prediction accuracy [25]. A hierarchical RNNs-based model was put out by Zhang et al. and employed an attention mechanism to map representations while combining short dependency path (SDP) and sentence sequence to gain semantic information [26,27]. A skeletal LSTM technique was developed by Jiang et al. to extract the internal structure of DDIs successfully [28].

Additionally, deep learning-based approaches, experts at extracting high-quality pharmacological characteristics, have seen extensive use in biology with encouraging outcomes [29,30]. To extract local and global properties of pharmaceuticals using the network, Karim et al. built a knowledge graph that considered the integration of CNN and LSTM models [31]. A factorization auto-encoder was used by Chu et al. to learn the representations of complicated nonlinear drug interactions [32]. To forecast DDIs, Liu et al. developed the multimodal deep auto-encoder known as DDI-MDAE based on shared latent representation [33]. Zhang et al. proposed a model DANN-DDI that predicted drug-drug interaction and associated events with it [34].

**Table S1.** Different body parts split. We use DrugBank (version 5.1.0) to select the body parts associated with the drugs' action and classify them. In the left column, the parts are mentioned, and the right column shows the number of drugs associated with each part.

| Body part split | Count of Node_id |
|---|---|
| Circulatory system | 119 |
| Digestive system | 76 |
| Endocrine system | 12 |
| Immune system | 26 |
| Integumentary system | 38 |
| Lymphatic system | 38 |
| Musculoskeletal system | 58 |
| Nervous system | 233 |
| Reproductive system | 60 |
| Respiratory system | 48 |
| Urinary system | 17 |
| Whole body | 116 |

**Table S2.** Drugs in different body-split. The DrugBank IDs are mentioned here for each body parts.

| Body-split | Drugs |
|---|---|
| Circulatory system | DB01098,DB00243,DB00390,DB11362,DB01118,DB00178,DB00436,DB00841,DB00264,DB01092,DB08819,DB00866,DB01136,DB00883,DB00898,DB01223,DB01167,DB01241,DB08816,DB01056,DB08907,DB01029,DB00727,DB09026,DB06209,DB09080,DB00594,DB08897,DB08931,DB00682,DB06712,DB00379,DB00612,DB00343,DB01426,DB01394,DB04846,DB00006,DB00353,DB01023,DB01244,DB00542,DB00211,DB00227,DB01396,DB00629,DB06268,DB00758,DB00270,DB00584,DB01253,DB00627,DB00946,DB08932,DB00661,DB01074,DB08860,DB01275,DB00945,DB00280,DB01412,DB00988,DB00204,DB01116,DB00640,DB00968,DB00528,DB00325,DB00180,DB00368,DB00175,DB01039,DB06217,DB01193,DB00820,DB04861,DB00908,DB00641,DB04855,DB01095,DB00598,DB00700,DB00805,DB01429,DB00790,DB00796,DB00519,DB00384,DB01195,DB00332,DB01197,DB01274,DB03756,DB00521,DB00177,DB01076,DB01054,DB01612,DB00381,DB00278,DB00869,DB00834,DB01035,DB00484,DB01020,DB01194,DB00206,DB01182,DB00808,DB06403,DB01240,DB00559,DB00401,DB00374,DB00421,DB01349,DB01228,DB04920,DB01021, |
| Digestive system | DB06689,DB01033,DB00432,DB01283,DB01233,DB00811,DB00592,DB00993,DB00585,DB01103,DB00836,DB00568,DB00673,DB09039,DB00889,DB00795,DB00850,DB00440,DB00903,DB06800,DB09049,DB02701,DB00737,DB00736,DB00602,DB01046,DB05521,DB09078,DB00927,DB00851,DB00501,DB00165,DB01184,DB00969,DB00366,DB00804,DB00433,DB01586,DB00759,DB00695,DB00134,DB00140,DB00863,DB00377,DB00613,DB05990,DB08896,DB01069,DB05351,DB00281,DB09462,DB01393,DB01176,DB00704,DB00213,DB00338,DB00508,DB01129,DB11691,DB00104,DB00604,DB08910,DB13155,DB09131,DB01014,DB06777,DB00806,DB00152,DB00183,DB00643,DB09280,DB08949,DB01115,DB08820,DB00322,DB00448, |
| Endocrine system | DB00451,DB05294,DB00550,DB00146,DB00763,DB00441,DB00910,DB01012,DB00389,DB06410,DB01101,DB01583, |
| Immune system | DB00451,DB05294,DB00550,DB00146,DB00763,DB00441,DB00910,DB01012,DB00389,DB06410,DB01101,DB01583,DB00283,DB00388,DB01075,DB00719,DB00471,DB00835,DB00455,DB00852,DB08906,DB01071,DB00748,DB00637,DB01222,DB00950,DB00967,DB09555,DB00557,DB01246,DB00427,DB00588,DB00915,DB00318,DB00615,DB11614,DB00537,DB04835, |

| | |
|---|---|
| Integumentary system | DB00549,DB00240,DB05219,DB00091,DB00769,DB06813,DB08828,DB00857,DB02300,DB01026,DB00916,DB00936,DB00337,DB08881,DB04794,DB08877,DB00887,DB01110,DB00799,DB00179,DB01013,DB00459,DB09031,DB01211,DB00845,DB01216,DB01179,DB00792,DB04930,DB00523,DB02709,DB01007,DB00724,DB00591,DB00199,DB01137,DB00755,DB06691, |
| Lymphatic system | DB01262,DB13874,DB00188,DB00309,DB04868,DB00563,DB00973,DB01254,DB09053,DB00444,DB00987,DB00694,DB00746,DB06616,DB00276,DB00997,DB06412,DB04898,DB00261,DB00115,DB01204,DB06228,DB00169,DB08901,DB08804,DB00884,DB01177,DB00307,DB01280,DB00541,DB06176,DB01169,DB00358,DB00928,DB00157,DB00158,DB12483,DB11581, |
| Musculoskeletal system | DB01219,DB01234,DB00741,DB00163,DB00749,DB00316,DB00443,DB00297,DB00860,DB01420,DB00720,DB01338,DB06736,DB00395,DB00172,DB06774,DB00788,DB01050,DB00608,DB00697,DB00356,DB00814,DB09120,DB01009,DB06725,DB01245,DB01138,DB00861,DB01173,DB01611,DB00469,DB00482,DB01166,DB01400,DB01097,DB00437,DB00500,DB00712,DB00554,DB08895,DB01384,DB01032,DB01685,DB01043,DB00586,DB00812,DB00328,DB00991,DB01144,DB00461,DB05676,DB01380,DB00605,DB00877,DB01296,DB00924,DB00153,DB06756, |
| Nervous system | DB00753,DB01085,DB00972,DB00683,DB00699,DB01576,DB08918,DB01171,DB01183,DB00819,DB00252,DB01558,DB01122,DB09396,DB00186,DB00347,DB00446,DB00575,DB06201,DB00752,DB00195,DB00543,DB01623,DB03128,DB01175,DB01149,DB01621,DB00818,DB00822,DB00696,DB00320,DB01186,DB00656,DB01224,DB01117,DB00360,DB04841,DB00966,DB08922,DB06016,DB00371,DB00344,DB00876,DB00458,DB00315,DB05381,DB01161,DB00849,DB01544,DB00933,DB00545,DB09118,DB00962,DB00540,DB00327,DB05541,DB00708,DB00231,DB00906,DB00571,DB00831,DB01355,DB00402,DB00425,DB09128,DB09148,DB04953,DB00477,DB00690,DB06413,DB01142,DB00476,DB00393,DB00921,DB01577,DB00934,DB00998,DB00633,DB00870,DB01238,DB00674,DB00312,DB00810,DB08811,DB00334,DB06144,DB11611,DB01192,DB01198,DB00751,DB01215,DB00205,DB00411,DB00740,DB09166,DB00324,DB00949,DB01589,DB00953,DB00420,DB00313,DB00989,DB01587,DB01559,DB00182,DB00768,DB06216,DB00228,DB01174,DB08880,DB00794,DB00434,DB00321,DB01100,DB00527,DB00176,DB00208,DB00599,DB00593,DB04844,DB09068,DB09119,DB06237,DB00202,DB01011,DB01156,DB04946,DB00754,DB03147,DB00268,DB06148,DB00472,DB00216,DB06637,DB01018,DB00745,DB00897,DB00996,DB00289,DB00617,DB00842,DB01107,DB00952,DB06654,DB05271,DB01065,DB01151,DB01104,DB00122,DB00185,DB00628,DB00323,DB00502,DB08883,DB01588,DB00721,DB00292,DB00679,DB00373,DB00363,DB00490,DB00703,DB01037,DB01154,DB04786,DB00918,DB01119,DB00829,DB01068,DB01320,DB00409,DB00623,DB00711,DB00979,DB01242,DB01159,DB00776,DB08815,DB01236,DB01595,DB00780,DB00273,DB00555,DB00802,DB01403,DB00622,DB06701,DB09167,DB00143,DB01580,DB00589,DB01624,DB05246,DB00564,DB06210,DB00285,DB00161,DB00370,DB06700,DB06262,DB01221,DB06204,DB00875,DB00422,DB00262,DB00726,DB06702,DB00392,DB00669,DB01437,DB01143,DB00843,DB00494,DB00513,DB00391,DB01337,DB00518,DB00546,DB00418,DB01235,DB00714,DB00184,DB00980,DB08868,DB06684,DB01064,DB00907,DB00246,DB00677,DB00909,DB00734,DB01367,DB08893, |
| Reproductive system | DB00346,DB00367,DB12332,DB09262,DB01036,DB01059,DB00457,DB00496,DB00957,DB01395,DB00203,DB06267,DB09074,DB09389,DB06207,DB00590,DB00624,DB08899,DB04908,DB00715,DB00882,DB00570,DB04839,DB00858,DB06772,DB00717,DB00867,DB09123,DB06789,DB01062,DB01128,DB00499,DB00977,DB00467,DB00209,DB00294,DB01591,DB00255,DB01431,DB06713,DB00215,DB00655,DB00784,DB01357,DB04574,DB04938,DB00917,DB00862,DB00783,D |

| | |
|---|---|
| | B08867,DB00603,DB01196,DB06738,DB05812,DB01708,DB00665,DB01126,DB01229,DB00515,DB00248, |
| Respiratory system | DB09076,DB00871,DB06282,DB00201,DB01418,DB01303,DB01410,DB12267,DB00956,DB01045,DB08911,DB00764,DB08912,DB01114,DB00233,DB00277,DB00394,DB09063,DB00514,DB00773,DB00254,DB01407,DB01466,DB09082,DB00317,DB00397,DB00951,DB01408,DB00938,DB06605,DB00778,DB01364,DB00744,DB09330,DB06739,DB06716,DB01435,DB01001,DB12364,DB00983,DB00642,DB01411,DB01135,DB01409,DB00572,DB08865,DB13139,DB00435, |
| Urinary system | DB00960,DB05039,DB00398,DB00035,DB06589,DB01165,DB06212,DB00925,DB00648,DB00136,DB01268,DB00357,DB00150,DB00706,DB06287,DB08875,DB00396, |
| Whole body | DB01083,DB04871,DB05239,DB12001,DB09130,DB00914,DB00668,DB01217,DB00170,DB01212,DB06757,DB00266,DB01351,DB00295,DB00495,DB01191,DB00130,DB01015,DB00242,DB06695,DB00678,DB00412,DB06335,DB00636,DB00162,DB06626,DB00631,DB11828,DB00990,DB01259,DB00978,DB00912,DB00245,DB00688,DB01124,DB06710,DB00191,DB00947,DB06155,DB01248,DB00865,DB11793,DB00445,DB01041,DB01006,DB00222,DB01024,DB00544,DB00454,DB01227,DB09143,DB09048,DB00984,DB01252,DB00959,DB00127,DB00731,DB00675,DB00872,DB00539,DB00620,DB01628,DB03615,DB06203,DB00428,DB00121,DB01094,DB01005,DB08439,DB04335,DB01120,DB08882,DB09073,DB00813,DB06594,DB11737,DB01105,DB00619,DB00470,DB01016,DB00487,DB01067,DB00635,DB06595,DB00904,DB00672,DB00131,DB01392,DB00963,DB01132,DB00481,DB01261,DB04845,DB00497,DB00361,DB00757,DB01028,DB00530,DB00491,DB01406,DB01131,DB00647,DB00173,DB08827,DB01590,DB01200,DB01213,DB00290,DB00257,DB00333,DB00118,DB00468,DB00847,DB00193,DB00166,DB09198, |

**Table S3.** Performance of our model in different negative sample sized datasets.

| Sample size | Precision | Recall | F1 | Accuracy |
|---|---|---|---|---|
| 1:1 | **0.99** | **0.99** | **0.99** | **0.99** |
| 1:2 | 0.98 | **0.99** | 0.98 | 0.99 |
| 1:3 | 0.48 | **0.99** | 0.64 | 0.73 |
| 1:4 | 0.50 | **0.99** | 0.64 | 0.77 |
| 1:5 | 0.50 | **0.99** | 0.67 | 0.77 |

**Table S4.** Effect of GraphSAGE layers on the performance of our model. We use a set of layers {1,2,3,4,5} for our analysis, and the experiment is done on the body-part split (endocrine system).

| No of layer | Precision | Recall | F1 | Accuracy |
|---|---|---|---|---|
| 1 | 0.96 | 0.50 | 0.66 | 0.74 |
| 2 | **0.98** | **0.85** | **0.91** | **0.92** |
| 3 | 0.93 | 0.53 | 0.65 | 0.73 |
| 4 | 0.94 | 0.51 | 0.66 | 0.74 |
| 5 | 0.91 | 0.52 | 0.64 | 0.72 |

**Table S5.** Effect of batch size on the performance of our model. We use a set of {16, 32, 64, 128, 256, 512} dimension batch sizes for our analysis, and the experiment is done on the body-part split (endocrine system).

| Batch size | Precision | Recall | F1 | Accuracy |
|---|---|---|---|---|
| 16 | 0.92 | 0.50 | 0.65 | 0.72 |

| | | | | |
|---|---|---|---|---|
| 32 | 0.93 | 0.51 | 0.66 | 0.73 |
| 64 | 0.95 | 0.50 | 0.65 | 0.73 |
| 128 | 0.96 | 0.50 | 0.65 | 0.73 |
| 256 | **0.98** | **0.85** | **0.91** | **0.92** |
| 512 | **0.98** | 0.50 | 0.66 | 0.74 |

**Table S6.** Effect of learning rates on the performance of our model. We use a set of learning rates {0.1,0.01,0.001,0.0001} for our analysis, and the experiment is done on the body-part split (endocrine system).

| **Learning rate** | **Precision** | **Recall** | **F1** | **Accuracy** |
|---|---|---|---|---|
| 0.1 | 0.80 | 0.50 | 0.62 | 0.70 |
| 0.01 | 0.83 | 0.53 | 0.65 | 0.77 |
| 0.001 | **0.98** | **0.85** | **0.91** | **0.92** |
| 0.0001 | 0.95 | 0.52 | 0.67 | 0.75 |

**Table S7.** Training time analysis between UniGEN-DDI and DANN-DDI.

| **Training data** | **Time (Hours) for UniGEN-DDI** | **Time (Hours) for DANN-DDI** |
|---|---|---|
| 10% | 0.74 | 1.42 |
| 20% | 1.47 | 2.71 |
| 30% | 2.20 | 3.75 |
| 40% | 2.93 | 4.79 |
| 50% | 3.67 | 6.04 |
| 60% | 4.44 | 7.25 |
| 70% | 5.18 | 8.34 |
| 80% | 5.95 | 9,62 |
| 90% | 6.69 | 10.75 |
| 100% | 7.51 | 11.92 |

**Table S8.** Comparison of body-part split performance of UniGEN-DDI with DANN-DDI and DPDDI. The number of drugs associated with each part is shown in the bracket.

| **Body Split** | **Metrics** | **DDIMDAE** | **RANEDDI** | **DPDDI** | **DANNDDI** | **UniGEN-DDI** |
|---|---|---|---|---|---|---|
| Circulatory system (119) | Precision | 0.88 | 0.94 | 0.93 | 0.97 | 0.97 |
| | Recall | 0.49 | 0.50 | 0.58 | 0.77 | 0.71 |
| | F1 | 0.63 | 0.65 | 0.71 | 0.86 | 0.82 |
| | Accuracy | 0.63 | 0.78 | 0.77 | 0.87 | 0.85 |
| Digestive system (76) | Precision | 0.86 | 0.95 | 0.94 | 0.98 | 0.98 |
| | Recall | 0.44 | 0.50 | 0.52 | 0.84 | 0.82 |
| | F1 | 0.58 | 0.66 | 0.67 | 0.90 | 0.89 |
| | Accuracy | 0.71 | 0.73 | 0.73 | 0.91 | 0.90 |
| Urinary system (17) | Precision | 0.84 | 0.94 | 0.95 | 0.98 | 0.94 |
| | Recall | 0.51 | 0.54 | 0.54 | 0.83 | 0.83 |
| | F1 | 0.63 | 0.68 | 0.69 | 0.90 | 0.88 |
| | Accuracy | 0.71 | 0.77 | 0.75 | 0.90 | 0.91 |
| Endocrine system (12) | Precision | 0.83 | 0.94 | 0.95 | 0.97 | 0.98 |
| | Recall | 0.45 | 0.52 | 0.55 | 0.84 | 0.85 |
| | F1 | 0.58 | 0.67 | 0.70 | 0.90 | 0.91 |

| | | | | | | |
|---|---|---|---|---|---|---|
| | Accuracy | 0.67 | 0.77 | 0.75 | 0.91 | 0.92 |
| Immune system (26) | Precision | 0.86 | 0.92 | 0.93 | 0.98 | 0.98 |
| | Recall | 0.51 | 0.55 | 0.57 | 0.92 | 0.91 |
| | F1 | 0.64 | 0.69 | 0.71 | 0.95 | 0.94 |
| | Accuracy | 0.65 | 0.78 | 0.76 | 0.95 | 0.95 |
| Lymphatic system (38) | Precision | 0.83 | 0.94 | 0.96 | 0.98 | 0.98 |
| | Recall | 0.49 | 0.48 | 0.53 | 0.87 | 0.81 |
| | F1 | 0.62 | 0.64 | 0.68 | 0.92 | 0.89 |
| | Accuracy | 0.65 | 0.70 | 0.70 | 0.92 | 0.89 |
| Musculoskeletal system (58) | Precision | 0.86 | 0.95 | 0.95 | 0.98 | 0.97 |
| | Recall | 0.51 | 0.52 | 0.56 | 0.91 | 0.81 |
| | F1 | 0.64 | 0.67 | 0.70 | 0.93 | 0.89 |
| | Accuracy | 0.65 | 0.69 | 0.76 | 0.94 | 0.89 |
| Reproductive system (60) | Precision | 0.84 | 0.93 | 0.96 | 0.98 | 0.97 |
| | Recall | 0.51 | 0.52 | 0.55 | 0.85 | 0.81 |
| | F1 | 0.63 | 0.67 | 0.70 | 0.90 | 0.88 |
| | Accuracy | 0.71 | 0.70 | 0.75 | 0.91 | 0.89 |
| Integumentary system (38) | Precision | 0.86 | 0.92 | 0.96 | 0.97 | 0.96 |
| | Recall | 0.44 | 0.51 | 0.50 | 0.84 | 0.58 |
| | F1 | 0.58 | 0.66 | 0.66 | 0.90 | 0.72 |
| | Accuracy | 0.71 | 0.74 | 0.77 | 0.92 | 0.78 |
| Respiratory system (48) | Precision | 0.84 | 0.93 | 0.94 | 0.98 | 0.96 |
| | Recall | 0.51 | 0.51 | 0.58 | 0.88 | 0.57 |
| | F1 | 0.63 | 0.66 | 0.72 | 0.93 | 0.71 |
| | Accuracy | 0.71 | 0.71 | 0.74 | 0.94 | 0.77 |

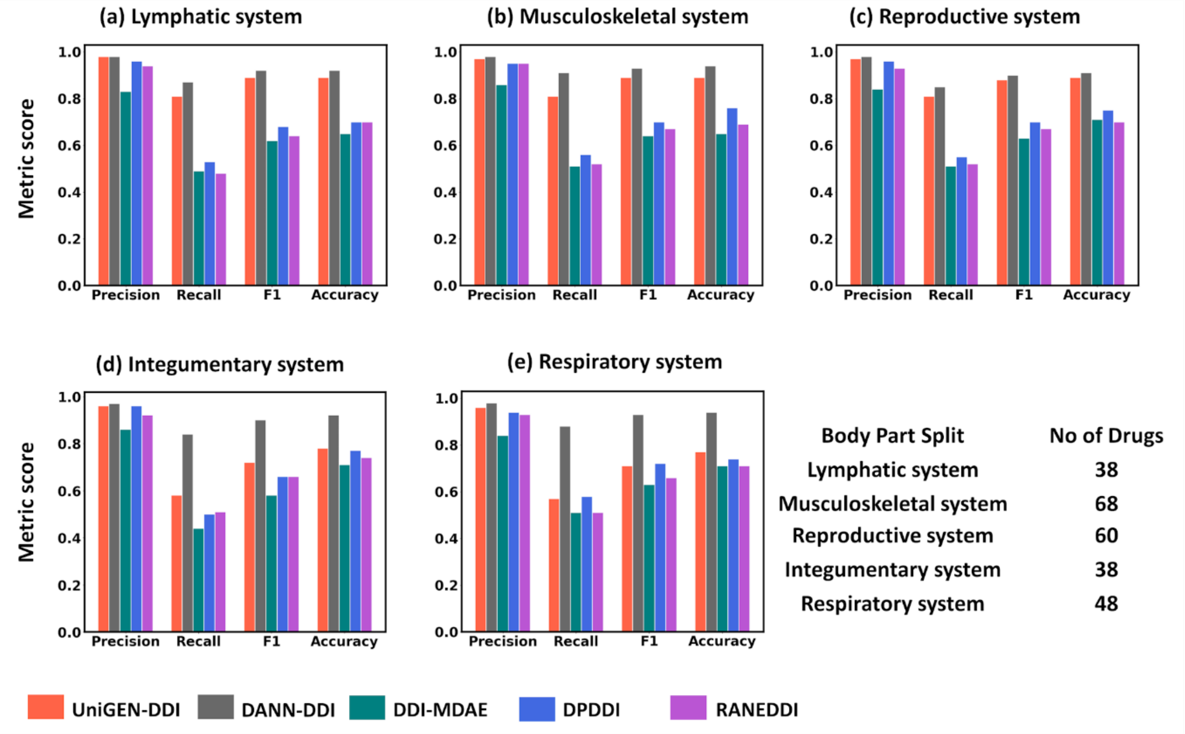


**Figure S1.** Comparison of UniGEN-DDI with state-of-the-art models in different body part splits.
